# EPHA2 Regulates SOX2 during Esophageal Development

**DOI:** 10.1101/2024.10.08.617209

**Authors:** Tianxia Li, Yosuke Mitani, Ricardo Cruz-Acuña, Tatiana A. Karaksheva, Varun Sahu, Cecilia Martin, Hiroshi Nakagawa, Joel Gabre

## Abstract

The human esophagus, derived from the anterior foregut endoderm, requires proper dorsal-ventral patterning for development. The transcription factor SOX2, crucial in this process, when dysregulated, leads to congenital esophageal abnormalities. EPHA2, a receptor tyrosine kinase, is vital in various developmental processes and cancer models, where it activates SOX2. This study demonstrates that EPHA2 regulates SOX2 expression during esophageal development using human iPSCs and iPSC-derived human esophageal organoids (HEO). Inhibition of EPHA2 decreased iPSC-derived HEO formation and SOX2 expression. These findings provide evidence of EPHA2 as being a key regulator of SOX2 signaling in early esophageal development.

**Graphical Abstract:** 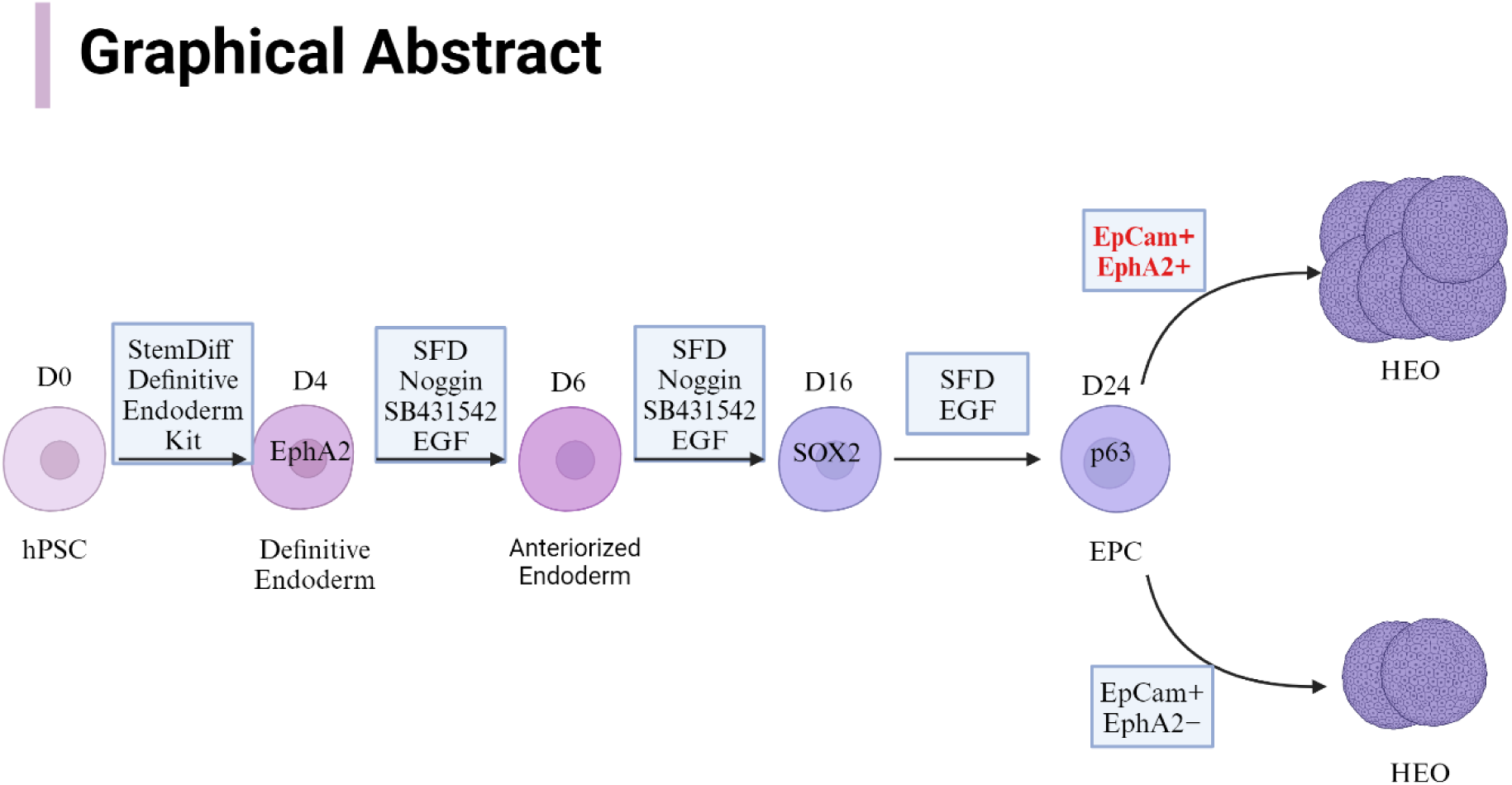
SFD: Serum-Free Differentiation media; EPC: esophageal progenitor cells; HEO: human esophageal organoids Created with BioRender.com

**Highlights:**

- SOX2 is crucial for proper esophageal development.
- EPHA2 is a receptor tyrosine kinase involved in various developmental processes.
- EPHA2 activates SOX2.
- Inhibition of EPHA2 decreased SOX2 expression and human esophageal organoid formation.

## Introduction

The human esophagus is lined by a non-keratinized stratified squamous epithelium that arises from the anterior foregut endoderm (AFE), which emerges around 4 weeks of development (Zhang *et al*. 2021). AFE then gives rise to the esophagus from the dorsal portion of the anterior foregut while the ventral anterior foregut gives rise to the trachea and lung (Zhang *et al* 2017). The SOX2 transcriptional factor is expressed dorsally while the NKX2.1 transcriptional factor is expressed ventrally, and they mediate these patterning steps. In human diseases and mouse models, the failure of SOX2 expression during the dorsal-ventral patterning step leads to congenital abnormalities of the esophagus, such as esophageal atresia and trachea-esophageal fistula (Que *et al.* 2007; Williamson *et al.* 2006). The Bone Morphogenic Proteins (BMPs) are known negative regulators of SOX2 (Trisno *et al.* 2018). Noggin, an inhibitor of BMP, is preferentially expressed along the dorsal aspect of the AFE, thereby aiding in SOX2 expression and patterning (Que *et al.* 2006). Positive regulators promoting SOX2 expression and esophageal development during this pattern step require identification and elucidation.

EPHA2 is a receptor tyrosine kinase important in many biological processes. It belongs to the EPH receptor family of receptor tyrosine kinases with an extracellular ligand-binding domain and an intracellular domain containing tyrosine residues, which upon phosphorylation, can regulate several signaling pathways. In early development, EPHA2 regulates differentiation of embryonic stem cells (Fernandez-Alonso *et al.* 2020). During organ development, EPHA2 is important in regulating eye lens development, branching morphogenesis of the kidney and mammary glands (Vaught *et al*. 2009; Park *et al.* 2013). In cancer models, EPHA2 is known to activate SOX2 to promote cancer cell stemness (Liang *et al*. 2023). In human esophageal development, SOX2 is known to mark the early esophageal basal cell compartment (Que *et al*. 2007). It is unknown if EPHA2 also marks the early fetal esophagus and regulates SOX2. Herein, we demonstrate using human fetal esophageal tissues, human induced pluripotent stem cells (iPSC) and iPSC-derived human esophageal organoids (HEO) that EPHA2 marks the early esophageal basal cell compartment and precedes SOX2 expression during AFE patterning. Functionally, we demonstrate that RNA interference mediated knockdown of EPHA2 impairs iPSC-derived HEO formation with decrease in SOX2 expression.

## Results and Discussion

### Expression of EPHA2 during Human Esophageal Development

To delineate the expression of EPHA2 and SOX2 during esophageal development, we examined their expression in human fetal esophageal tissues. We performed immunofluorescence staining of human fetal esophageal tissues at 7, 10, 14, and 17 weeks, respectively (Fig. 1 and Sup Fig. S1). Tracheo-esophageal separation occurs between weeks 4-8 of development (Que *et al*. 2007). At 7 weeks of gestation, the esophageal epithelium is pseudostratified columnar with limited SOX2 expression. However, at 10 weeks of gestation, esophageal epithelial cells start to express EPHA2, SOX2 and p63. After 14 weeks of gestation, cells with increased EPHA2, SOX2 and p63 expression align to form the basal cell layer. Overlying differentiated epithelial cells exhibit decreased expression of these proteins. The expression of Ki-67, indicating cell proliferation, is primarily observed in the basal layer and the layer just above it. These findings demonstrate co-localization of EPHA2, SOX2 and p63 during early esophageal development with further restriction to the basal compartment during differentiation.

**Figure 1.**
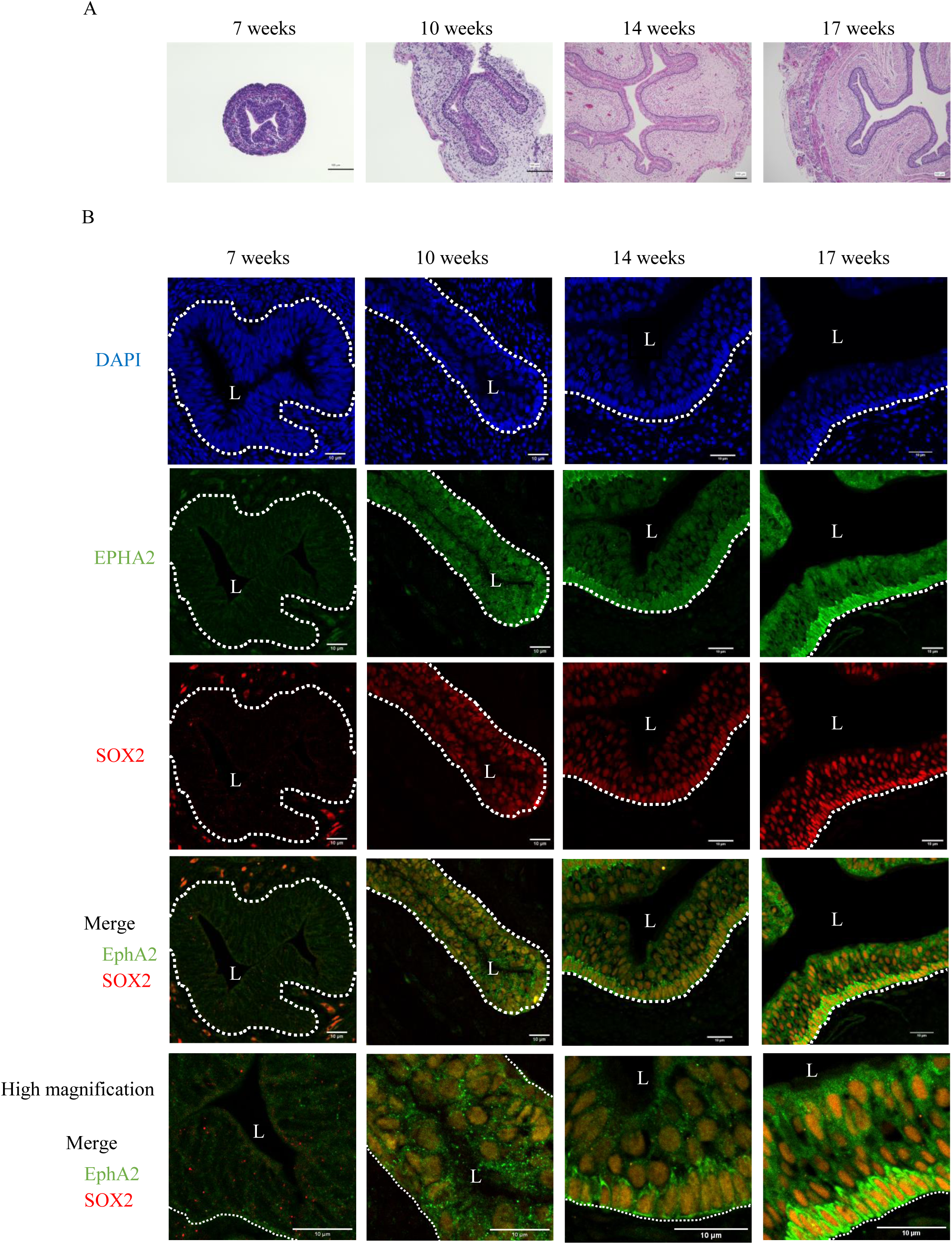
Development of the human esophagus and expression of EPHA2 and SOX2. (*A*) H&E staining of the human fetal esophagus at various gestational weeks. Scale bar indicates 100µm. (*B*) Immunofluorescence staining of the human fetal esophagus at various gestational weeks. Scale bar indicates 10µm. The white dotted line indicates the basal cell layer. L: esophageal lumen.

### EPHA2 is expressed during the differentiation of human pluripotent stem cells into esophageal basal cells

To dissect the role of EPHA2 during esophageal development, we performed *in vitro* differentiation of BU3 NGST human iPSCs (Boston University) into esophageal basal cells (Jacob *et al.,* 2017, Zhang *et al*. 2018). We collected total RNA and protein at differentiation days D0, D4, D6, D16 and D24 during esophageal basal cell differentiation. Day 6 represents development of the AFE based on previous studies using this model system (Zhang *et al*. 2018). Day 24 represents the early esophageal basal cell. Both EPHA2 mRNA and protein levels increased at D4 followed by increased SOX2 mRNA and protein levels at Day 6 (Fig. 2A, B), which represents the AFE that has been patterned dorsally. We observed an increase of p63 following SOX2 induction during the final stages of differentiation into the esophageal basal cell by day 24 consistent with what has been reported previously about p63 in this context (Watanabe *et al.* 2014). These findings demonstrate that during differentiation of iPSCs into early esophageal basal cells, EPHA2 expression precedes SOX2 expression.

**Figure 2.**
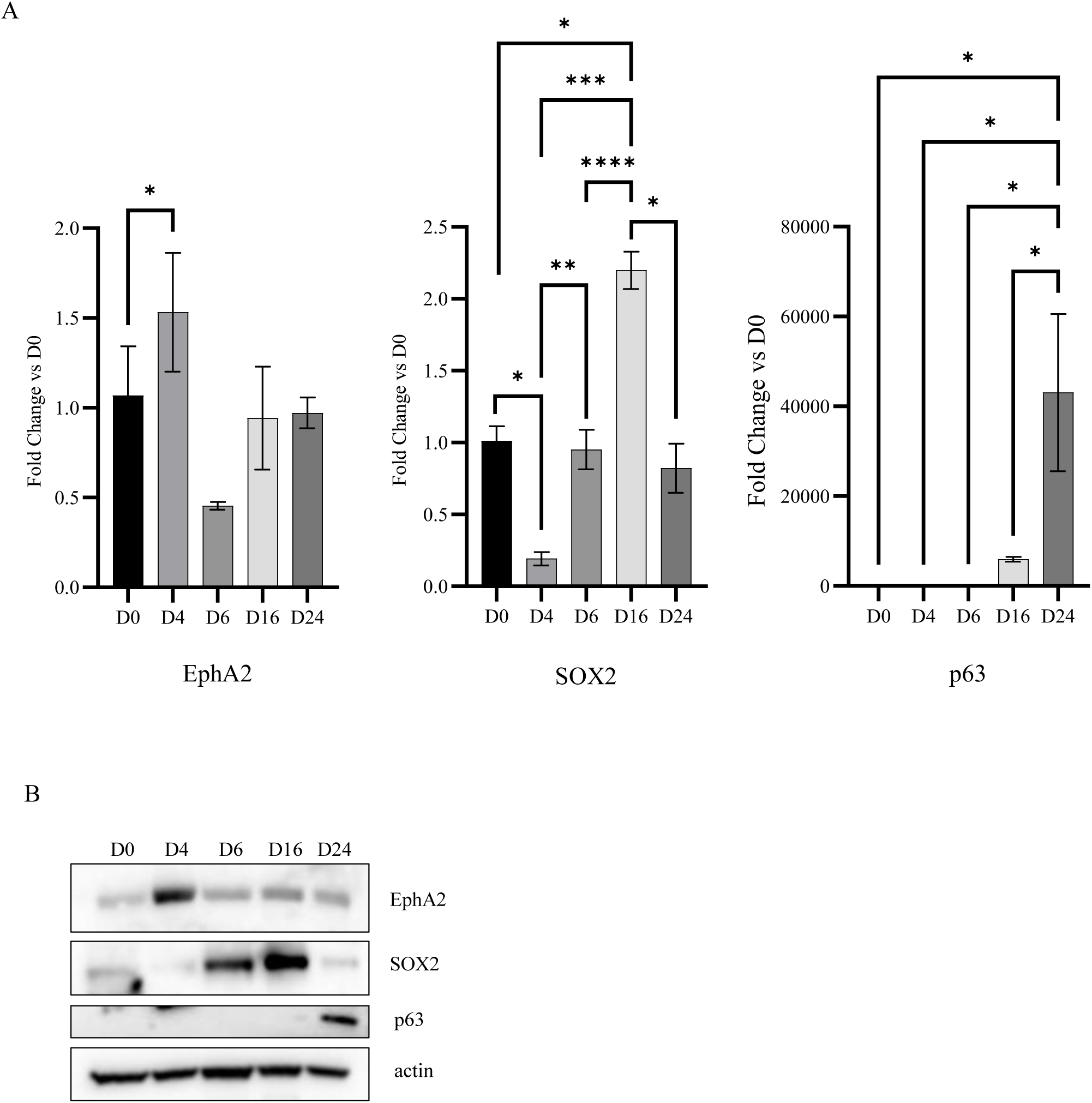
EPHA2 is expressed in the differentiation of Human Pluripotent stem cells into esophagus epithelial cells. *(A)* Representative qRT-PCR of EPHA2, SOX2 and p63 expression of a time course of a differentiation of BU NGST iPSC cells into esophagus epithelial cells, with fold change (2-ΔΔCt) ± SD, n=3, One-way ANOVA, (*) *P* ≤ 0.05; (****) *P* ≤ 0.0001, Error bars represent SD. (*B*) Immunoblots of EPHA2, SOX2 and p63 of a time course of the differentiation of BU3 NGST PSCs, actin is used as a loading control.

### Inhibition of EPHA2 decreases iPSC-derived Human Esophageal Organoid (HEO) formation and SOX2 expression

Esophageal basal cells derived from iPSCs were cultured to form 3D organoids as previously described (Zhang *et al.* 2021). To determine function of EPHA2 inhibition in HEO formation, we selected two established pharmacological inhibitors: NVP-BHG712, which is a pan-inhibitor of Eph receptor tyrosine kinases (Zhao *et al.* 2024, Martiny-Baron *et al.* 2010) and ALW II-41-27, which is a potent and more specific inhibitor of EphA2 (Zhang *et al.* 2019, Choi *et al.* 2009), and performed 3D CellTiter-Glo cell viability assays and organoid formation assays (Fig. 3A, B). The cell viability assay showed ALW II-41-27 had a stronger effect at a lower dose of 300nM than NVP-BHG712, which revealed a similar effect but only at a dose previously reported toxic dose (10µM) (Martiny-Baron *et al.* 2010). Similarly, ALW II-41-27 showed 50% inhibition of HEO formation at 200nM as did NVP-BHG712, but at the higher dose of 2µM (Fig. 3C, D). These results indicate that specific inhibition of EPHA2 may impair HEO formation and viability. Furthermore, to investigate the target efficacy of EPHA2 inhibitor, we examined phosphorylation of both EPHA2 and one of its downstream targets, STAT3, by immunoblotting. ALW II-41-27 demonstrated the best suppression of pEPHA2 (Ser897), pSTAT3 (Tyr705) and SOX2. Taken together, these results demonstrate that EPHA2 inhibition can impair HEO formation, likely through decreased STAT3 activation and SOX2 expression.

**Figure 3.**
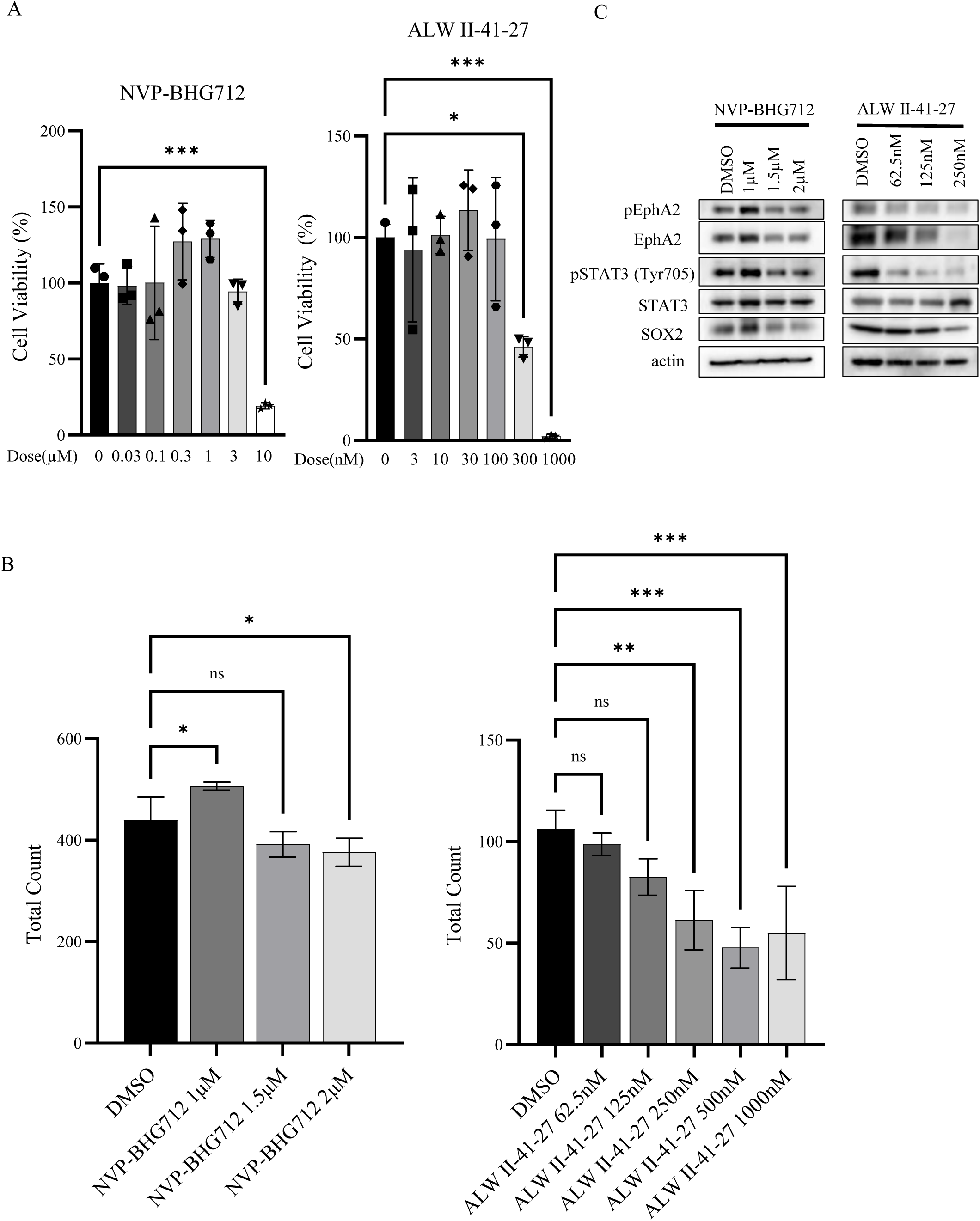
Block EPHA2 by NVB-BHG712 and ALW II-41-27 show inhibition of organoids formation. After flow sorting for Epcam, all Epcam positive cells are counted and resuspended in matrigel. 20K cells are seeded in 50ul matrigel/well in 24-well plate. Cells are grown in Matrigel in organoid culture media for seven days before treatment. (*A*) Cell viability (n=3) after 72-hour treatment with NVP-BHG712 or ALW II-41-27 at indicated doses. Cell viability is normalized to DMSO control group and expressed as a percentage of maximum viability. (*) *P* ≤ 0.05; (***) *P* ≤ 0.001, Error bars represent SD. (*B*) Total organoids are counted by Celigo imagine system after 5 days treatment with NVP-BHG712 or ALW II-41-27 at indicated doses (refresh treatment every two days). n=3, One-way ANOVA, (*) *P* ≤ 0.05; (**) *P* ≤ 0.01; (***) *P* ≤ 0.001, Error bars represent SD. (*C*) Representative immunoblots of phosphor-EPHA2 (Ser897), EPHA2, phosphor-STAT3 (Tyr705), STAT3 and SOX2 in HEOs after 5 days treatment (refresh treatment every two days) with NVP-BHG712 or ALW II-41-27 at indicated doses. DMSO is used as vehicle control, actin is used as a loading control.

### Knockdown of EPHA2 decreases human esophageal organoid (HEO) formation

To validate the function of EPHA2 in HEOs, we constructed three doxycycline-inducible EPHA2 shRNA iPSC cell lines (parental line BU3 NGST) for differentiation into HEOs, aiming to verify the contribution of EPHA2 to HEO formation via SOX2 signaling. Doxycycline induction of EPHA2 shRNA (#1) in iPSC cells resulted in the most efficient knockdown of EPHA2 expression (Fig. 4A). Given this result, we selected shRNA#1 for further differentiation experiments. EPHA2, SOX2 and p63 were expressed sequentially from D0 to D24 during the development of HEO in our shRNA#1 knockdown cell line (Fig. 4B,C), similarly as the wild-type (WT) iPSC line undergoing differentiation (Fig 2). Next, we evaluated HEO formation. Total HEO counts were decreased significantly upon doxycycline mediated induction of EPHA2 shRNA (Fig. 4D). We evaluated the expression of pEPHA2 (Ser897) and EPHA2 at protein level (Fig. 4E) and mRNA level (Fig. 4F) upon EPHA2 depletion, which reduced further pSTAT3 (Tyr705) and SOX2 (Fig 4E). In summary, these results indicate that EPHA2 knockdown leads to decreased HEO formation and decreased SOX2, expression highlighting that EPHA2 may be upstream of SOX2 signaling during HEO development.

**Figure 4.**
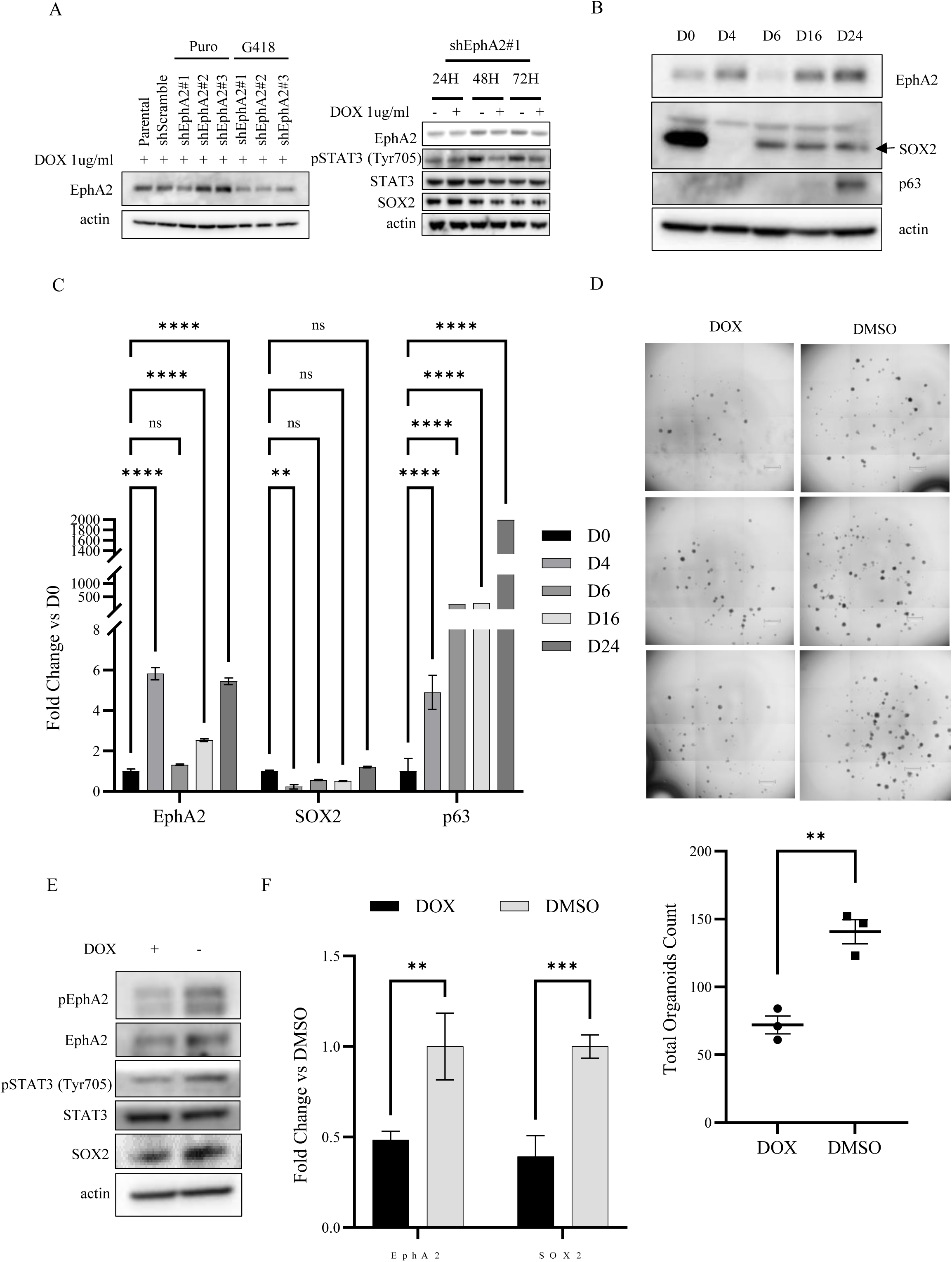
Knockdown of EPHA2 decreases human esophagus organoids formation. (*A, left panel*) Representative immunoblots of EPHA2 after Doxycycline 1µg/mL 72 hours in parental BU3 NGST PSCs, scramble shRNA and three inducible EPHA2 shRNA stable BU3 NGST PSCs in two different backbones, actin is used as a loading control. (*A, right panel*) Immunoblots of EPHA2, phosphor-STAT3 (Tyr705), STAT3 and SOX2 in inducible EPHA2 shRNA#1 PSCs with Doxycycline induced 24h, 48h and 72h, DMSO is used as vehicle control, actin is used as a loading control. (*B*) Representative immunoblots of EPHA2, SOX2 and p63 of a time course of the differentiation of inducible EPHA2 shRNA#1 BU3 NGST PSCs, actin is used as a loading control. (*C*) Representative qRT-PCR of EPHA2, SOX2 and p63 expression of a time course of a differentiation of inducible EPHA2 shRNA#1 BU NGST PSCs into esophagus epithelial cells, with fold change (2-ΔΔCt) ± SD, n=3, One-way ANOVA, (**) *P* ≤ 0.01; (****) *P* ≤ 0.0001, Error bars represent SD. (*D*) Bright field Images of organoids formation in 24 well plates. Organoids are shown as black circles and size variety. Total number of organoids each well are counted by Celigo image system. n=3, unpaired t test, (**) *P* ≤ 0.01, Error bars represent SD. (*E, F*) EPHA2 depletion with shRNA#1 is validated by qRT-PCR (*F*) with fold change (2-ΔΔCt) ± SD, n=3, two-way ANOVA, (**) *P* ≤ 0.01; (***) *P* ≤ 0.0005, error bars represent SD and immunoblots of phosphor EPHA2 (Ser897) and EPHA2 (*E*) after 5 days Doxycycline (1ug/mL) introduction in HEOs. Immunoblots of phosphor STAT3 (Tyr705), STAT3 and SOX2 are decreased after EPHA2 was depleted.

### Knocking down EPHA2 disrupts the maintenance of an undifferentiated basal cell state

To investigate the impact of EPHA2 on HEOs, we conducted bulk RNA sequencing and analysis comparing EPHA2 knockdown HEOs to WT HEOs. Inducible EPHA2 shRNA-expressing HEOs were seeded and treated with either doxycycline or DMSO starting from Day 7, with RNA collected on Day 14. Bulk RNA sequencing was performed to identify differentially expressed genes, which were visualized using a volcano plot (Figure 5A). Knockdown of EPHA2 resulted in significant upregulation of 17 genes and significant downregulation of 20 genes, with the upregulated genes including esophageal differentiation markers, such as KRT13 and KRT16. Next we conducted GO analysis for each set of upregulated and downregulated genes (Figure 5B, C). The upregulated gene set included genes involved in epithelial cell differentiation and keratinization, as shown in the Biological Process section of Figure 5B. These findings suggest that knockdown of EPHA2 causes HEOs to lose their ability to maintain an undifferentiated esophageal basal cell state, but switch towards a differentiation state.

**Figure 5.**
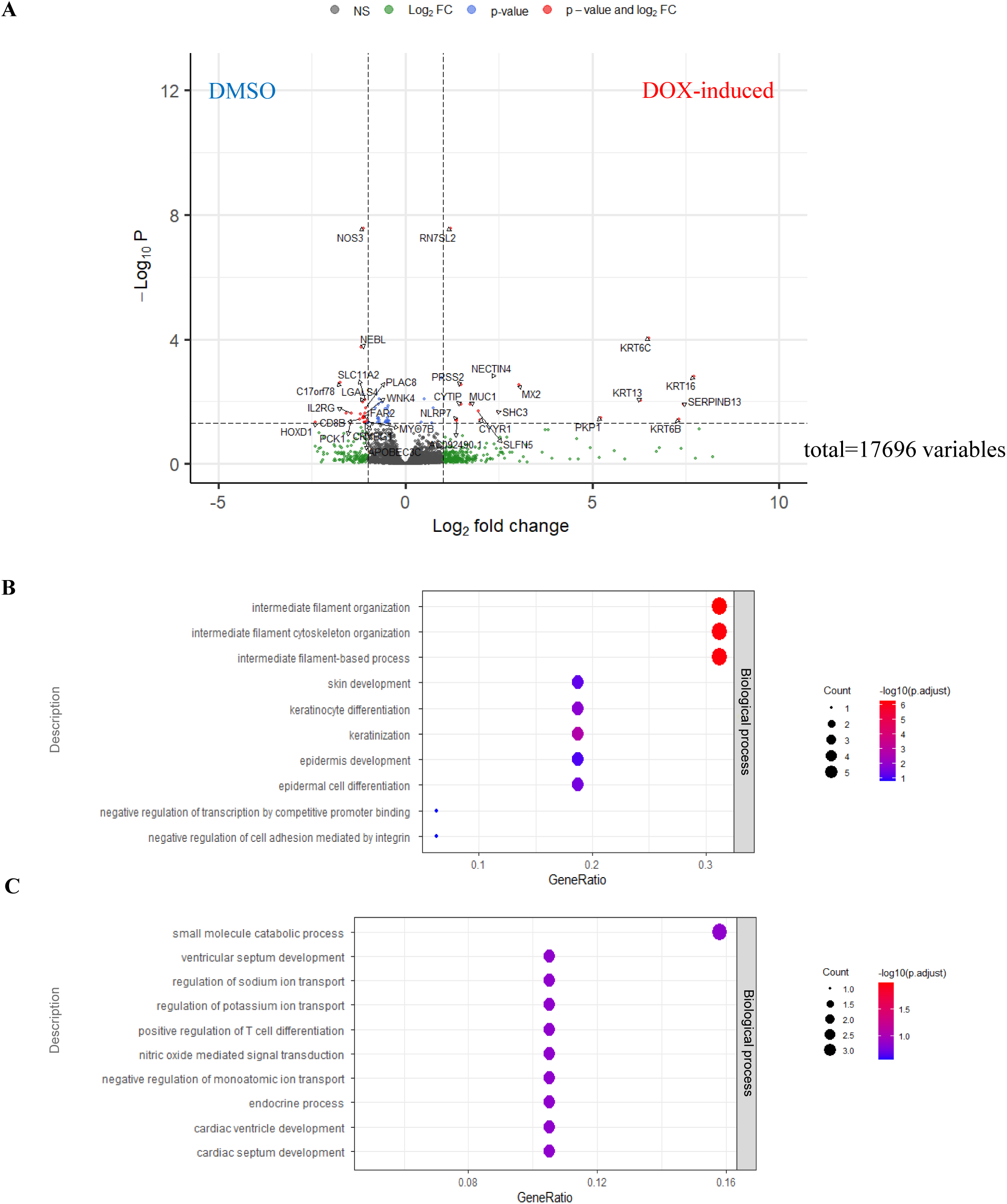
Identification and functional analysis of differentially expressed genes (DEG) following EPHA2 knockdown. (*A*) DEGs are shown in a Volcano plot comparing the DOX and DMSO groups, with red dots indicating significant DEGs where the Log2fold change is greater than 1 and the adjusted p-value is less than 0.05. (*B*) Upregulated DEGs and (*C*) downregulated DEGs were analyzed using the biological process category of Gene Ontology (GO). The plots show GO terms with the size of each plot representing the gene counts and the color indicating the significance of the enrichment.

In summary, SOX2 is a transcription factor critical to a squamous epithelial cell’s fate, as it is important in patterning the anterior foregut endoderm towards an esophageal cell linage. Additionally, SOX2, along with p63, is a marker of the esophageal basal cells. During development, BMP factors are well known to be important factors in inhibiting SOX2 expression during the AFE patterning step. Noggin, expressed dorsally in the AFE, is known to be an inhibitor of BMP signaling. It remains unknown if there are positive regulators of SOX2 during this patterning step. In our study we have identified that EPHA2 expression precedes SOX2 expression in AFE patterning and EPHA2 is co-expressed with SOX2 in the early esophageal basal cell. Utilizing human iPSCs and iPSC HEOs we have identified that knockdown of EPHA2 decreases HEO formation and SOX2 expression using an inhibitor of and inducible shRNA knockdown of EPHA2. Furthermore, RNA sequencing analysis suggested that suppression of EPHA2 plays a significant role in the maintenance of the undifferentiated basal cell state. Recently, a study involved the generation of iPSCs from three patients with type C esophageal atresia/tracheo-esophageal fistula without other malformations compared to healthy controls (Raad *et al*. 2022). The authors found transient SOX2 dysregulation during anterior foregut patterning. Our studies provide evidence that EPHA2 may be upstream of SOX2 and possibly contributing to this dysregulation. Understanding this signaling axis could provide further mechanistic insights into the regulation of SOX2 and may contribute to the understanding of congenital diseases of the esophagus, including esophageal atresia and tracheoesophageal fistula in sporadic cases.

## Materials and Methods

### Cell culture

BU3 NGST iPSC line (Jacob *et al.,* 2017) was purchased from Boston University and was cultured in mTeSR Plus media (Stemcell Technologies) with 1% penicillin/streptomycin (Gibco) in feeder free conditions on 6 well plates coated with 1:100 Matrigel (BD Bioscience, San Jose, CA) in 1X PBS. iPSCs were grown at 37°C with 5% CO2 with media changed every second day. Cells were passaged using Dispase (Corning) when > 70% confluent.

### Esophageal Epithelial Basal Cell Differentiation from iPSCs

Differentiation into esophageal basal cells was performed according to previously reported methods (Zhang *et al*. 2021). To generate endoderm, BU3 NGST iPSCs were detached using Accutase (Stemcell Technologies) and cultured in mTeSR Plus medium (Stemcell Technologies) with 10 μM Y-27632 (Selleckchem) on 6-well plates coated with 1:100 diluted Matrigel (Corning). After 24 hours (Day 1), the medium was replaced with Definitive Endoderm medium plus Supplement MR and CJ (Stemcell Technologies). After another 24 hours (Day 2), the cells were incubated in Definitive Endoderm medium plus Supplement CJ. Serum-Free Differentiation (SFD) medium was prepared as follows: 375 ml of reconstituted Iscove’s DMEM (Corning), 125 ml of Ham’s F-12 (Corning), 5 ml of N2 (Thermo Fisher Scientific), 10 ml of B27 (Thermo Fisher Scientific), 3.3 ml of 7.5% Bovine Albumin Fraction V Solution (Thermo Fisher Scientific), 19.6 μl of 11.5M monothioglycerol (Sigma-Aldrich), 500 μl of L-Ascorbic Acid (Sigma-Aldrich), and 1 ml of 50 mg/ml Primocin (Invitrogen). On Day 4, the cells were passaged into new Matrigel-coated 6-well plates and anterior foregut progenitor cells were further induced by culturing endoderm in the Serum Free Differentiation (SFD) medium plus 10 μM SB431542 (Selleckchem), 200 ng/ml Noggin (R&D Systems), and 100 ng/ml EGF (Peprotech), with 10 μM Y-27632 for 24 hours. The SFD medium plus 10 μM SB431542, 200 ng/ml Noggin, and 100 ng/ml EGF was then used for another 48 hours (Days 5-6). Cells were maintained at 5% O2 / 90% N2 / 5% CO2 from Days 1-6. To induce esophageal basal cell differentiation, anterior foregut progenitor cells were cultured from Days 7 to 15 in the SFD medium plus 10 μM SB431542, 200 ng/ml Noggin, and 100 ng/ml EGF. From Days 16 to 24, cells were maintained in the SFD medium plus 100 ng/ml EGF. Cells were cultured at 95% air / 5% CO2 from Day 7 onwards. On Day 24, the cells were sorted using FACS as follows. For cell surface marker staining, cells were dissociated with Accutase plus DNase (Thermo Fisher Scientific) and 10 μM Y-27632 and stained with PE/Cyanine7 anti-human CD326 (EpCAM) antibody (BioLegend) in FACS buffer (1× PBS, 2% FBS, plus DNase and 10 μM Y-27632) for 1 hour with live/dead staining dye (SYTOX, Thermo Fisher Scientific) to exclude dead cells. Stained cells were analyzed using a BD FACSAria II (BD Biosciences) and EPCAM-positive and GFP-negative cells were sorted. The iPSC-derived esophagus epithelial cells were cultured in 3D organoid culture as described below.

### 3D Organoid Culture of the iPSC-derived esophageal basal cells

The iPSC-derived esophageal basal cells were counted and seeded at 20,000 cells per 50 μl Matrigel (Corning) per well of a 24-well plate, and medium was added after the Matrigel solidified. The organoid culture medium included the SFD medium supplemented with 10 μM Y-27632, 10 μM SB431542, 10 μM CHIR99021, 20 ng/ml FGF2, and 200 ng/ml EGF. Media was changed every other day. Bright-field imaging of organoids and measurement and quantification of the number and size of organoids were performed using the Celigo Image Cytometer (Nexcelom) and its analytic algorithms.

### Small molecular inhibitors and treatment

NVB-BHG712 and ALW II-41-27 were purchased from Selleck Chemicals and dissolved in DMSO before use. After flow sorting EpCAM positive cells, 20,000 cells were resuspended in 50ul of Matrigel in 24-well plates in triplicate. After organoids solidified, 500ul of HEO media was applied and replaced every 2 days. Organoids were grown 7 days before drug treatment. On day 7, organoids were exposed to DMSO or indicated inhibitors.

### shRNAs and cloning

shRNA oligos (all primer sequences used were listed in Table 1) were ordered from Sigma MISSION® shRNA and cloned into Tet-pLKO-puro (Addgene 21915) and Tet-pLKO-neo (Addgene 21916). Tet-pLKO-puro-Scrambled (Addgene 47541) was used as control. Detailed cloning procedure was performed as a previously described (Wiederschain *et al* 2009).

**Table 1.**
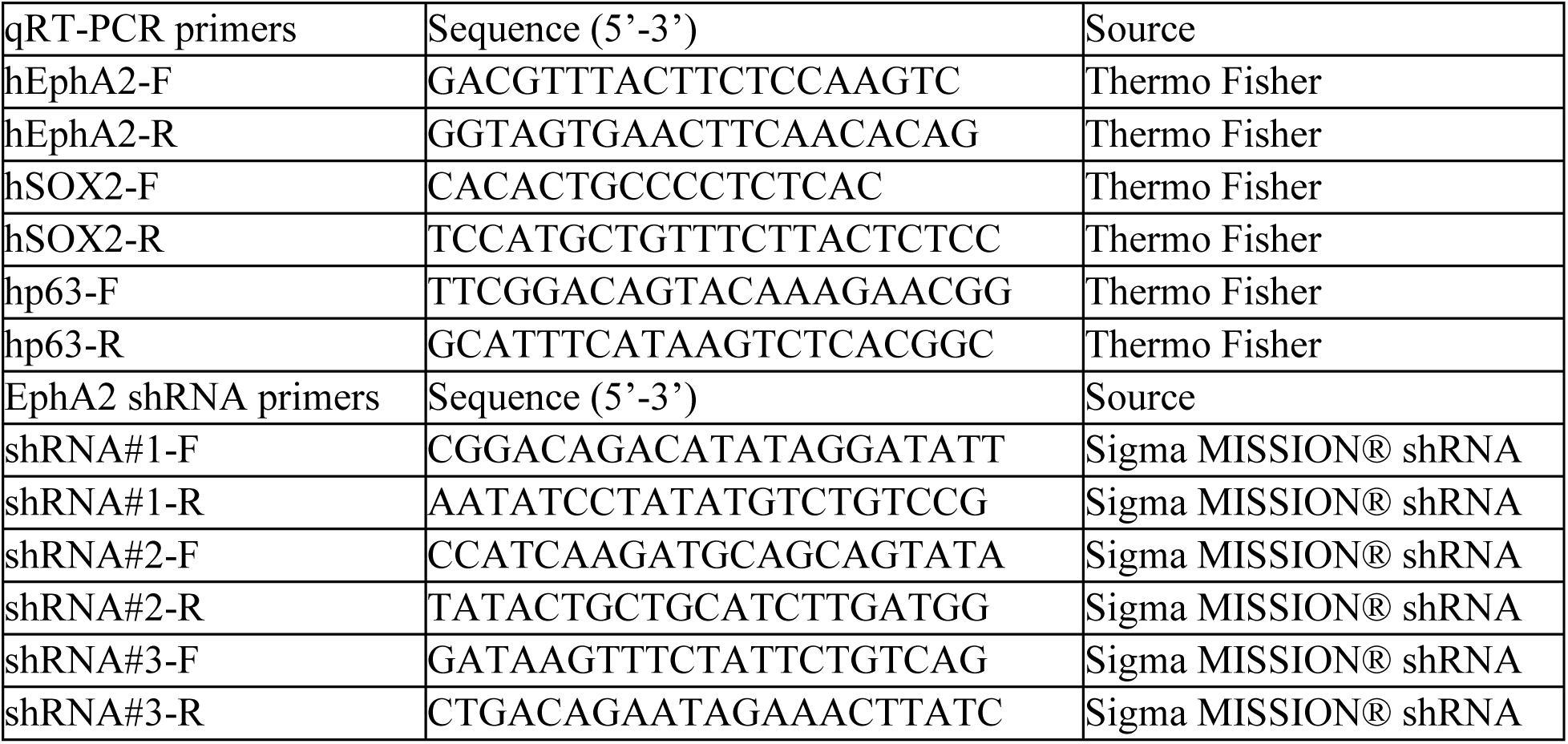

### Lentiviral Production

Lentiviral production was performed using HEK293T cells as described on the Broad Institute Genetic Perturbation Platform (GPP) portal: http://portals.broadinstitue.org/gpp/public/....lots).

Briefly, lentiviral particles were produced by transfecting HEK293T cells with psPAX2 (Addgene 12260) and pMD2.G (Addgene 12259) plasmids using lipofectamine 2000 (ThermoFisher). Media was changed 16 hours following transfection. Lentivirus-containing media were collected after 24 hours and 48 hours.

### Cell Viability Assay

After flow sorting EpCAM positive cells, 1000 cells were resuspended in 25ul of Matrigel in 96- well plates in triplicate. After organoids solidified, 100ul of HEOs media was applied and replaced every 2 days. Organoids were grown 7 days before drug treatment. On day 7, organoids were exposed to DMSO or indicated inhibitors for 72 hours. Cell viability was quantified by using 3D CellTiter-Glo Luminescent Cell Viability assay (Promega G9682) according to manufacturer’s instructions.

### Western blot

iPSCs were collected at different stages during differentiation and placed in 1X protein lysate buffer which contains RIPA buffer (Boston Bioproducts BP-115x), protease inhibitor (Sigma Aldrich) and phosphatase inhibitor cocktail (Roch) then lysed on ice. For HEO collection, Matrigel surrounding the organoids was dissolved using TrypLE Solution (Corning) in a thermomixer at 37°C shaking at 800rpm for 5 minutes. The released organoids were pelleted and lysed in 1X protein lysate buffer. Immunoblot analysis was performed as previously described (Wong *et al*. 2018). Briefly, primary antibodies to pEphA2 (Ser897) (1:1000; Cell Signaling Technology 6347), EphA2 (1:1000; Cell Signaling Technology 6697), pSTAT3 (Tyr705) (1:1000; Cell Signaling Technology 9145) , STAT3 (1:1000; Cell Signaling Technology 12640), SOX2 (1:1000; Cell Signaling Technology 3579), p63 (1:1000; Cell Signaling Technology 13109), and actin (1:5000; Sigma A3853) were incubated overnight at 4C. Membranes were then washed three times for five minutes in TBS-T and were incubated with secondary antibodies (Thermo Fisher Scientific) diluted in blocking buffer for 1 hour. Amersham ECL Prime chemiluminescent detection reagent (GE Healthcare Life Sciences) was used to visualize protein expression. All immunoblots were conducted twice. Shown are representative results from one experiment.

### RNA isolation and qRT-PCR

Total RNA from iPSCs or HEO pellets was extracted with RNAqueous Micro Kit (Thermo Fisher) according to the manufacturer’s instructions. cDNA was synthesized by using iScript reverse transcription supermix for RT-PCR (Bio-Rad 1708841). qRT-PCR was performed with the StepOnePlus real-time PCR system (Applied Biosystems) by using Power SYBR Green PCR Master Mix (Applied Biosystems). All primer sequences used are listed in Table 1.

### RNA Sequencing and Enrichment Analysis

RNA samples were shipped on dry ice to Azenta (Burlington) for RNA sequencing. RNA-seq libraries were prepared using the NEBNext Ultra RNA Library Prep Kit (New England Biolabs) following the manufacturer’s instructions. Sequencing was performed on an Illumina NovaSeq (San Diego), generating paired-end reads. The raw sequencing data were processed and analyzed using Azenta’s standard bioinformatics pipeline, which includes quality control, read alignment to the reference genome, transcript assembly, and differential expression analysis. Differentially expressed genes were identified using the DESeq2 package. To interpret the biological significance of the differentially expressed genes, Gene Ontology (GO) enrichment analysis was performed using the clusterProfiler package in R (version 4.4.0).

### Immunofluorescence and Confocal Microscopy

Formalin-fixed paraffin-embedded samples were sectioned into 5 µm thick slices and stained with hematoxylin-eosin or for immunofluorescence. For immunofluorescence, the slides were subjected to antigen retrieval with 10 mM citrate buffer (pH 6.0) and stained with the following primary antibodies for 10 hours at 4 °C: mouse anti-EphA2 (1:50; Santa Cruz, sc-398832) and rat anti-SOX2 (1:200; Invitrogen, 14-9811-82). Subsequently, the slides were stained with the following secondary antibodies for 1 hour at room temperature: goat anti-mouse IgG AlexaFluor 488-conjugated (1:200; Invitrogen, A11001) and donkey anti-rat IgG AlexaFluor 555-conjugated (1:200; Invitrogen, A21434). Nuclei were stained with DAPI in the mounting medium, VECTASHIELD HardSet (Vector Laboratories). Images were obtained with a Nikon A1 confocal microscope (Nikon).

### Statistics analysis

Statistics analyses were performed using Prism, version 10.1.2. All data are presented as mean ± SD unless otherwise noted and analyzed by unpaired t-test, one-way ANOVA or two-way ANOVA. P values are denoted in figures by asterisks (\**P* ≤ 0.05, ** *P* ≤ 0.01, \*\*\**P* ≤ 0.001, \*\*\*\**P* ≤ 0.0001).

### Author Contributions

T.L., Y.M., and J.G. performed iPSC differentiation and organoid experiments. R.C.A. and V.S. provided technical assistances with experiments. T.L., Y.M., and J.G. conceptualized and designed studies. T.K., H.N. and C.M. assisted with data interpretation. T.L. Y.M., J.G. wrote the manuscript, and all authors provided revisions for the manuscript. All authors have read and agreed to the published version of the manuscript.

## Data Availability Statement

The original contributions presented in the study are included in the article/supplementary material, further inquiries can be directed to the corresponding author.

## Acknowledgments

This work was supported by the Herbert Irving Comprehensive Cancer Center Molecular Pathology Shared Resource, Organoid & Cell Culture Core of the NIH P30 P30DK132710 Columbia University Digestive and Liver Diseases Research Center, The Columbia Stem Cell Initiative’s Stem Cell Core.

